# A population-based human enCYCLOPedia for circadian medicine

**DOI:** 10.1101/301580

**Authors:** Marc D. Ruben, Gang Wu, David F. Smith, Robert E. Schmidt, Lauren J. Francey, Ron C. Anafi, John B. Hogenesch

## Abstract

The discovery that half of the mammalian protein-coding genome is clock-regulated has clear implications for medicine. Indeed, recent studies demonstrate time-of-day impact on therapeutic outcomes in human heart disease and cancer. Yet biological time is rarely given clinical consideration. A key barrier is the absence of information on the *what* and *where* of molecular rhythms in the human body. Here, we have applied CYCLOPS, an algorithm designed to reconstruct sample order in the absence of time-of-day information, to the GTEx collection of 632 human donors contributing 4,292 RNA-seq samples from 13 distinct human tissue types. We identify rhythms in expression across the body that persist at the population-level. This includes a set of ‘ubiquitous cyclers’ comprised of well-established circadian clock factors but also many genes without prior circadian context. Among thousands of tissue-divergent rhythms, we discover a set of genes robustly oscillating in cardiovascular tissue, including key drug targets relevant to heart disease. These results also have implications for genetic studies where circadian variability may have masked genetic influence. It is our hope that the human enCYCLOPedia helps drive the translation of circadian biology into prospective clinical trials in cardiology and many other therapeutic areas.

**One Sentence Summary:** Bioinformatic analyses on thousands of human tissue samples reveals an enCYCLOPedia of rhythmic gene expression across the body and identifies key translational opportunities for circadian medicine in cardiovascular disease.

## Introduction

Vital metabolic, endocrine, and behavioral functions demonstrate daily variation in humans *(1)* orchestrated by endogenous circadian clocks that drive tissue-specific rhythms in gene expression *(2)*. The discovery that ~ 50% of mammalian genes are expressed with 24 h rhythms *(3*, *4)* has broad physiological and pharmacological implications. Circadian medicine aims to incorporate this information to develop safer and more efficacious therapeutics. Recent studies show that time-of-day is critical to outcomes in heart surgery *(5)* and cancer treatment *(6*, *7)*. Despite this, neither clinicians nor drug developers typically consider biological time in practice. A key reason for this is the lack of understanding of the mechanisms by which the circadian clock governs rhythms in physiology and pathophysiology. Human time-series data have not been feasible at the spatial and temporal resolution of animal models *(8*–*10)*.

Importantly, state-of-the-art algorithms *now* allow us to build a population-level human circadian atlas. We have applied CYCLOPS *(11)* — an algorithm designed to reconstruct sample order in the absence of time-of-day information — to the Genotype-Tissue Expression (GTEx) collection *(12)* of 4,292 postmortem RNA-seq samples from 632 donors across 13 distinct tissue types. We found extensive population-scale rhythms in expression, with ~50% of protein-coding genes cycling in at least one of the 13 tissues. We identified a set of genes that oscillate across the majority of tissues. These include well-established “core clock” genes and also many with little or no prior circadian context. Interestingly, known clock genes exhibit robust oscillations and evolutionarily conserved phase relationships (e.g., ROR target transcriptional activators *BMAL1*, *NPAS2*, and *CLOCK* peak antiphase to their E-box targets *NR1D1*, *NR1D2*, *PER3*) despite noisy human data. In contrast to studies in model organisms, these data are population-level and show that the oscillator functions robustly despite many inter-individual differences (e.g. age, sex, health status). The finding that half the human genome is subject to rhythmic regulation has implications for many fields of medicine, and also large-scale genetics studies where associations may have been missed because circadian variability wasn’t accounted for.

Although the core clock network is shared across organ systems, the clear majority of rhythmic expression was tissue-specific. Circadian medicine hinges on reliable prediction of drug-target spatiotemporal dynamics. Focusing on the cardiovascular system, we connect over 100 drugs to 136 genes oscillating robustly in at least one of: atrial chamber, aorta, coronary artery, or tibial artery. The human heart and vasculature demonstrate 24 h rhythms in physiology. For example, blood pressure (BP) is higher during wake than sleep *(13)*. Further, the cardiomyocyte molecular clock in mice regulates BP *(14)*. Antihypertensives are used to decrease awake or sleep BP and can restore asleep dipping, re-establishing diurnal rhythms within the cardiovascular system *(15*, *16)*. Yet, little is known about the underlying mechanisms. We identified several classes of drugs whose target genes are rhythmic in the cardiovascular system, including standard-of-care antihypertensives (i.e. Ca^2+^ channel- and ß-blockers). These represent new opportunities for timed dosage in pursuit of circadian medicine.

## Results

### Population-level rhythms in gene expression across human anatomy

From the full GTEx collection of RNA-sequenced human donor samples, we ordered 13 of 51 tissues with CYCLOPS (fig. S1A). These 13 tissues were chosen because i) they had at least 160 samples, ii) the molecular clock network was preserved *(17)*, and iii) the orderings reproduced evolutionarily conserved phase relationships between core clock genes. In total, our analyses considered 632 donors characterized by a mix of ages, sex, and health status (Table 1).

**Table 1.** GTEx donor demographics. The 632 donors analyzed were a mix of males and females from age 20 to 79 yrs at time of death. Death was classified on the 4-point Hardy Scale, as follows. *Fast:* death due to accident, blunt force trauma or suicide, terminal phase estimated at < 10 min; *Sudden*-*Natural:* fast death of natural causes, sudden unexpected deaths of people who had been reasonably healthy, after a terminal phase estimated at < 1 h (e.g., sudden death from a myocardial infarction); *Intermediate:* death after a terminal phase of 1 to 24 h, patients who were ill but death was unexpected; *Slow:* death after a long illness, with a terminal phase longer than 1 day (e.g., cancer or chronic pulmonary disease); *Ventilator:* all cases on a ventilator immediately before death. Additional inclusion criteria are in Methods.

| **Sex** | **Male** | **Female** |  |  |  |  |
| --- | --- | --- | --- | --- | --- | --- |
|  | 66% | 34% |  |  |  |  |
| **Age** | **20-29** | **30-39** | **40-49** | **50-59** | **60-69** | **70-79** |
|  | 8% | 7% | 17% | 34% | 32% | 3% |
| **Death** | **Fast** | **Intermediate** | **Slow** | **Sudden-Natural** | **Ventilator** | **N/A** |
|  | **4%** | **5%** | **10%** | **26%** | **53%** | **1%** |

Cosinor analysis on ordered samples from each tissue identified thousands of genes whose expression met criteria for rhythmicity (false discovery rate *(18)* (FDR) < 0.05, relative amplitude (rAmp) ≥ 0.1, and goodness-of-fit (RSQ) ≥ 0.1 (fig. S2A, Data file S1). We found that 7,486 distinct genes cycle in at least one of the 13 tissues (Fig. 1A). This represents 44% of total distinct genes considered for analysis (16,906), which were limited to the top 15K expressed from each tissue. While the vast majority of cycling genes are protein-coding, a small share in each tissue are noncoding elements, including antisense RNAs, miRNAs, and lincRNAs (fig. S2B, Data file S1).

**Fig. 1.**
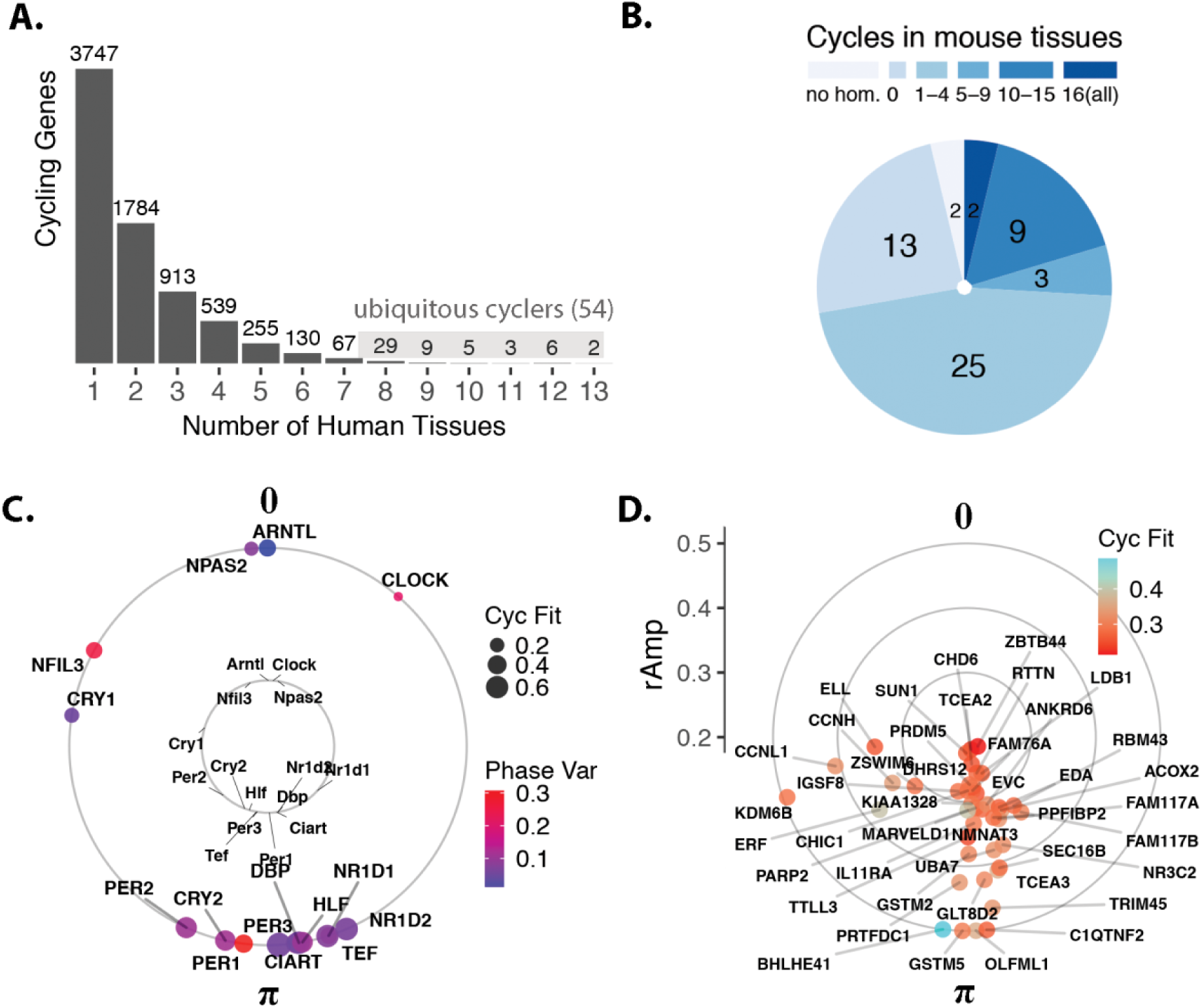
Population-level rhythms in gene expression across human anatomy. (**A**) Number of genes that met criteria for rhythmicity (FDR < 0.05, rAmp ≥ 0.1, RSQ ≥ 0.1) in at least one of the 13 tissues. Periodic analysis was performed by cosinor regression on expression values for 160 CYCLOPS-ordered samples per tissue, as discussed in Fig. S1 and Methods. See Fig. S2 for numbers of rhythmic genes discovered at varying FDR thresholds. (**B**) Set of 54 *ubiquitous* genes cycling in at least 8 of 13 human tissues, grouped by prior circadian context as reported in CircaDB (darkest blue: homologue cycles in 16/16 mouse tissues; lightest blue: not cycling in any of the 16 mouse tissues). (**C**) Average acrophases of core clock genes (external circle) across all human tissues compared to mouse (internal circle, average across 12 tissues reported in Zhang et. al.). Average cycling robustness (Cyc Fit) indicated by point size, and phase variability (Phase Var) indicated by color. Peak phase of *ARNTL* (human) or *Arntl* (mouse) was set as 0 for comparison. (**D**) Average acrophases of all other (non-core clock) human ubiquitous cycling genes. Genes located more distant from the center have larger average amplitudes of oscillation (rAmp) across tissues where they cycle.

We uncovered a set of 54 genes that cycle in 8 or more of the 13 tissues (Fig. 1A and B), referred to hereafter as ‘ubiquitous’. Among these are well-established clock and clock-controlled genes, that show phase correlations matching those observed in mouse — e.g., target transcriptional activators *BMAL1*, *NPAS2*, and *CLOCK* peak on average 12 h out of phase with their E-box targets *NR1D1*, *NR1D2*, and *PER3* (Fig. 1C). In addition to known clock factors, the ubiquitous set includes genes that do *not* cycle robustly in mice *(19)* (Fig. 1B and D, Data file S2).

### Robust oscillator and divergent output programs

Core clock genes ranked among the most robustly cycling genes across the majority of tissues analyzed (Fig. 2A). This aligns with observations from time-series studies — from invertebrates through non-human primates *(3*, *4*, *20)*. As the first large scale population analysis, we demonstrate that clock gene oscillations persist despite wide interindividual variability.

**Fig. 2.**
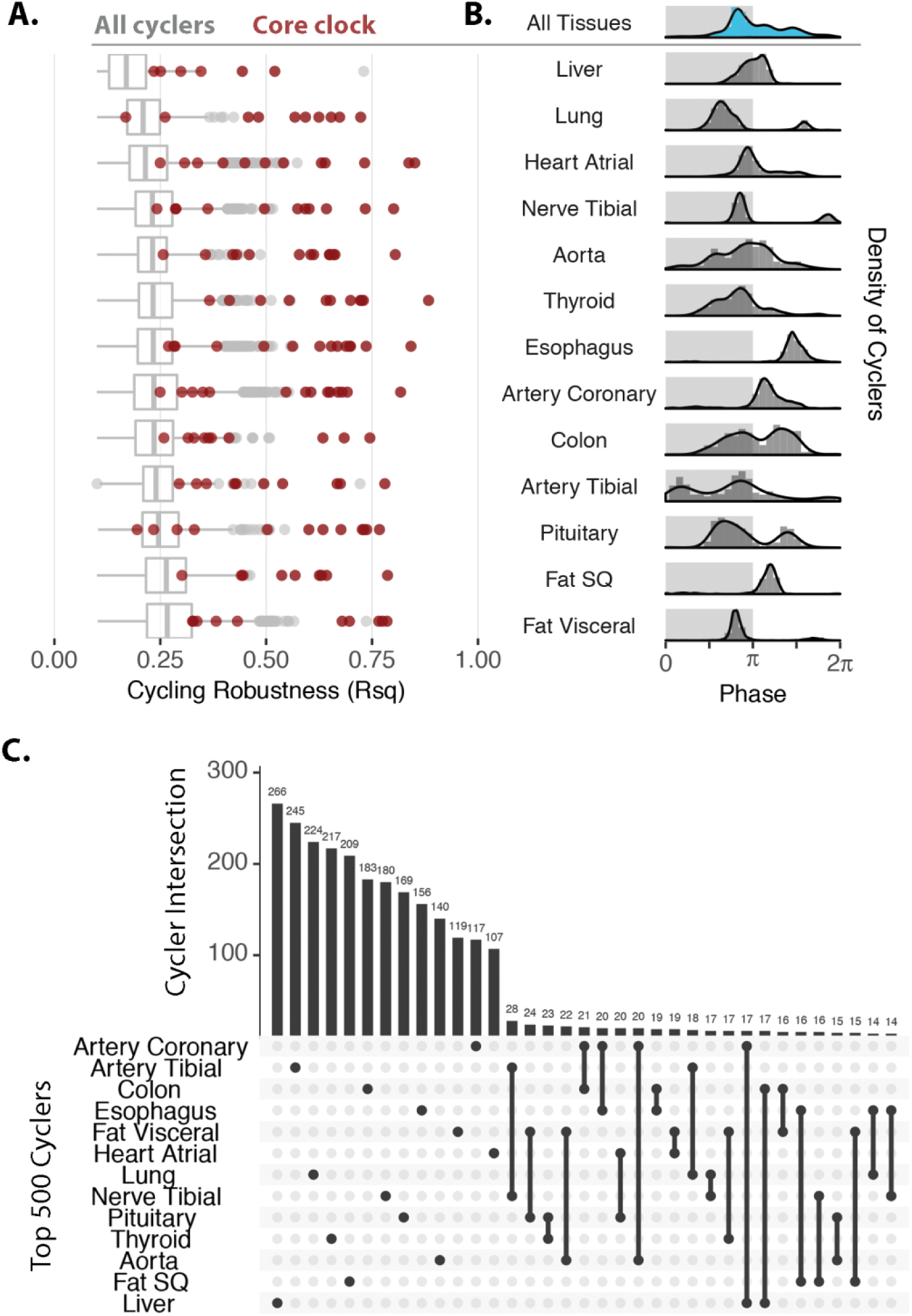
Robust tissue-specific output. (**A**) Distribution of robustness (RSQ, i.e., goodness of data fit to 24 h sine wave) for all cyclers (grey) in each tissue; clock genes (red) reside in the upper-quartile in each tissue. (**B**) Distribution of acrophases for cyclers differs between tissue types. Light and dark shading represents inferred active and inactive phase, based on *Arntl* expression which is known to peak in anticipation of the inactive period in mammals. (**C**) UpsetR *(23)* to rank and visualize the sizes of intersection between multiple sets (i.e., tissues). Restricting input to the top 500 cyclers by FDR from each tissue, the largest intersections describe genes whose expression cycles *only* in a single tissue.

While the core clock network is shared across tissues, overall transcriptional programs are not. The cycling transcriptomes of the 13 tissues differ in both phase distribution and genes (Fig. 2B and C). On the whole, most tissues display a prominent peak of rhythmic transcription at or near the end of the inferred inactivity phase, evident in single-tissue and composite (blue shaded) histograms (Fig. 2B). There are exceptions to this. Peak phases in esophagus, coronary artery, and subcutaneous fat occur later, during the inferred activity phase. Interestingly, and similar to observations from time-series profiles from murine models *(3)*, we detect strong unimodal or bimodal structures, depending on tissue type. While the physiological significance of this remains unknown, studies of heart rhythms, for example, suggest that these patterns reflect organ function *(21*, *22)*.

We applied UpsetR *(23)* to rank and visualize sizes of intersection (i.e. genes) between tissues (Fig. 2C). When restricting input to the top 500 cyclers (ranked by FDR) in each tissue we found that the largest 2-tissue intersection, between tibial artery and tibial nerve, is comprised of just 28 genes (5%) and has nearly expected overlap due to chance alone (Fisher’s exact test, p = 0.1). Indeed, most cycling genes belong to non-intersecting sets. For example, 266 of the top 500 liver cyclers (53%) are exclusive to liver. Anatomic divergence in clock output is a well characterized feature in model organisms, likely reflecting the nature of sampling across distinct organ systems. While precise mechanisms governing tissue specific circadian gene regulation require elaboration, its extent underscores the distinct functions of the human organs profiled.

Pathway analyses *(24)* identified evolutionarily conserved temporal regulation of numerous biological processes (fig. S3). Top enriched gene networks were tissue-specific, many with therapeutic relevance (e.g., ion transport in the heart and aorta), prompting us to look at drug targets.

### Dynamic drug targets

Previous analyses have linked circadian genes in mouse, and recently baboon, to specific drug entities *(3*, *4)*. Here, we sought to identify pharmacological connections to rhythms in the human population. Of the thousands of genes that cycle in at least one of the 13 tissues (fig. 2A), 917 (12%) of them encode proteins that either transport or metabolize drugs or are themselves drug targets (Fig. 3A) *(25)*. Overall, this connects thousands of different drugs — both approved and experimental — to nearly one thousand cycling genes. Figure 3B depicts genes that map to drugs with a defined mechanism for each tissue (either transport, metabolism, or drug action). Subsequent analyses focused on this set.

**Fig. 3.**
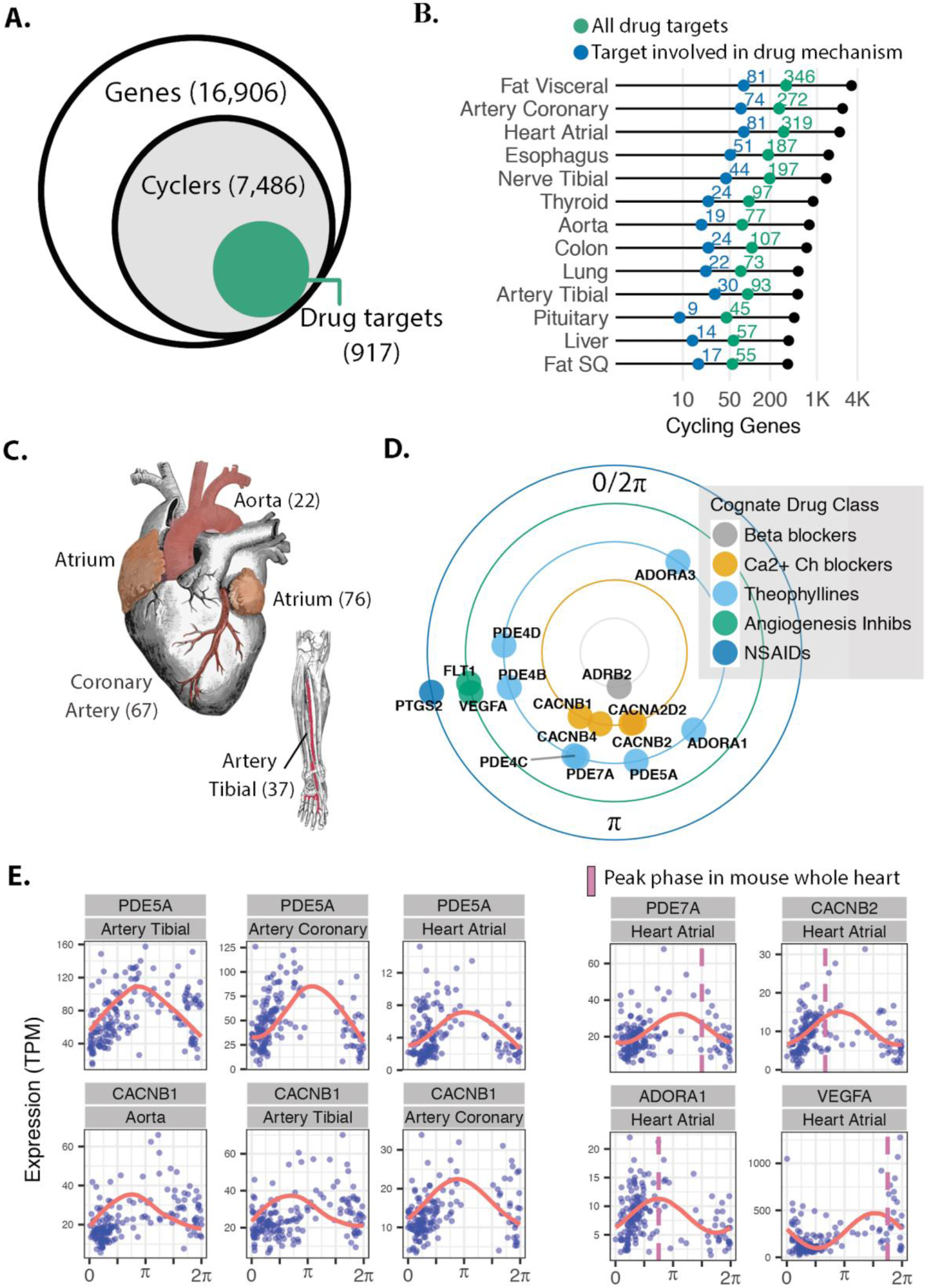
Pharmacological links to molecular rhythms in the human population. (**A**) Of 7,486 genes found to cycle in at least one of 13 human tissues sampled, 917 (12%) encode at least one drug target, transporter, or metabolizing enzyme (collectively referred to as ‘targets’). This represents a total of 2,764 drug entities, both approved and experimental as logged in Drugbank *(25)*, with targets that oscillate somewhere in the body. (**B**) Numbers of total (colored green) and pharmacologically active (colored blue) cycling drug targets by tissue type. (**C**) Cardiovascular-related tissues from ordered GTEx data. To enrich for high amplitude ‘cardio-cyclers’ we limited to genes in the upper 50^th^ percentile for relative amplitude within each tissue and FDR ≤ 0.1. Numbers of cardio-cyclers that meet these cycling criteria and are *also* drug targets are indicated in parentheses. (**D**) Cardio-cyclers targeted by select drug classes relevant to heart and vessel physiology. Plotted phase represents average peak phase across all cardiac tissues where that gene was found to cycle. (**E**) Expression values plotted as a function of sample phase for select cardio-cyclers. *Right*: genes that are also rhythmically-expressed in whole mouse heart, with peak phases indicated by pink dashed lines. *ARNTL* (human) or *Arntl* (mouse) was set to 0 for comparison.

Genes that cycle in human cardiovascular tissue (atrial chamber, aorta, coronary artery, or tibial artery) are targeted by many of these drugs. The cardiovascular system demonstrates 24 h rhythms in BP and contractility *(13)*. Mechanisms underlying daily variation in heart and peripheral vasculature, however, remain poorly understood. To reveal the strongest oscillating drug targets in cardiovascular tissue, we narrowed focus to the subset of ‘cardio-cyclers’ in the upper 50^th^ percentile for relative amplitude. This identified 136 drug targets cycling in at least one of the four tissues (Data file S3, Fig 3C, fig. S4). A large number of these have established relevance to cardiovascular physiology — many being key targets of standard-of-care agents in heart disease (Fig. 3D and E). For example, the family of calcium channel blockers, which inhibit Ca^2+^ influx into cells of the heart and blood vessels producing smooth muscle relaxation, target several rhythmically expressed Ca^2^+ channel subunits (*CACNA2D2*, *CACNB1*, *CACNB2*, *CACNB4*). Many of these agents are short-acting (half-lives < 6 h) and may be influenced by time of administration (Fig. 3D).

We identified a separate group of drug classes that, while not necessarily indicated for cardiovascular pathology, actively target this system (Fig. 3D and E). For example, theophyllines — bronchodilators prescribed for asthma and chronic obstructive pulmonary disease — function by inhibiting adenosine receptors (ADORA1/3) and multiple phosphodiesterases (PDEs) in the lung. These genes are also critical to normal heart function *(26*, *27)* and their pharmacologic inhibition can be cardiotoxic *(28)*. Additional examples include non-steroidal-anti-inflammatory drugs (NSAIDS) prescribed for pain, and anti-angiogenic agents commonly used in cancer, both associated with cardiovascular risk *(28*, *29)* and both with cardiac-cycling targets. Many targets also cycle in mouse heart or aorta with similar phase relationships (Fig. 3E, Data file S3) (whole heart and/or aorta, JTK Q-value ≤ 0.1) *(3*, *30*, *31)*, indicating evolutionary conservation of these targets and significance for application in circadian medicine.

## Discussion

Efforts toward precision medicine have largely centered around linking genetic variants to therapeutic response. Time-series profiles from mice *(3)*, and recently a nonhuman primate *(4)*, show extensive rhythms in transcript abundance and suggest that *circadian time* should factor into drug choice and use. However, while animal models were instrumental to our understanding of circadian biology, results from their study have had limited translational impact. Typically, animal model experiments use small numbers of animals of similar sex, age, and environmental conditions. These are critical variables in all clinical translation, including circadian medicine.

To address these limitations, we are moving to population based analyses. We applied CYCLOPS to thousands of unordered human samples taken from a heterogeneous population and found that nearly half of all analyzed genes are rhythmically expressed somewhere in the body. This approximates findings in mice, where the number of FDR-controlled cyclers discovered plateaus at ~50% of the protein-coding genome *(3)*. Nearly 1,000 of the human cyclers influence drug action. This far exceeds the number of human genes with allelic associations to a specific drug response *(25)*. A few drugs targeting several of these mechanisms are already taken with time-of-day dependence (e.g., steroids, short-acting statins). Nevertheless, this leaves many opportunities to explore time of dosage to improve the therapeutic index (i.e., promoting efficacy and limiting toxicity). Further, circadian variation has undoubtedly masked genetic influence. Development of algorithms to account for this variability may reveal new genetic associations — indeed the circadian clock may contribute to ‘the missing heritability’ problem.

Many drug metabolizing enzymes, transporters, and targets were found cycling in the cardiovascular system. Circadian clock-dependent variation in cardiac ion channels has been shown in animal models *(32*–*34)* and is proposed to underlie time-dependent arrhythmias and sudden cardiac death in humans *(35*, *36)*. We found that several L-type Ca^2+^ channel subunits — critical for cardiac polarization and targets of commonly prescribed antihypertensives — cycle in the human heart and vessels at population scale. Ca^2^+ channel blockers, however, are not routinely administered with regard to time of day. We propose that the kinetics of commonly used Ca^2^+ channel blockers (e.g. verapamil, nifedipine, and others) could be used to determine the most effective times of administration based on the windows of peak target abundance. In support of this notion, several papers report improved efficacy of nifedipine by timing dosage prior to bedtime (e.g., *(37)*).

The circadian system could also influence adverse events. Cardiotoxicity is the leading cause of drug discontinuation *(38)*. Drugs designed to target other organ systems can provoke adverse cardiac problems. For example, theophylline, commonly prescribed for pulmonary disease, can elicit heart palpitations that limit or preclude its use in some patients *(28).* We note that NSAIDS, typically prescribed for pain and inflammation, and anti-angiogenic agents typically prescribed for cancer, can cause cardiotoxic response *(29*, *39)*. Here we show that they also have cardio-cycling targets. For these agents, dosage time could be leveraged to avoid cardiotoxicity and perhaps retain efficacy.

Translational potential for the enCYCLOPedia hinges on a correspondence between transcript and protein. While prior work shows an estimated ~80% of rhythmic proteins in liver have corresponding mRNA rhythms *(40)*, the amplitudes and phases of target proteins will need to be determined empirically. An additional limitation of our findings is that inferred relationships between estimated transcript phase and external clock time (i.e. light:dark) relied upon the assumption that clocks in different tissues share a similar phase. While based on time-series studies of clock gene expression from multiple tissues in mouse (e.g. *BMAL1* peaks in anticipation of the inactive period), this assumption may not hold true at human population scale. To this point, Hughey et. al. show phase differences between central (i.e. brain) and peripheral (i.e. blood and skin) clocks *(41)*. The degree to which clocks in different organ systems are differentially-phased remains an outstanding question in circadian biology, but does not prevent application of circadian medicine.

Prospective trials designed to test the impact of time-of-day drug administration are required for clinical implementation. Wearable devices with continuous monitoring are being used to study a wide range of physiologic measures with daily rhythms *(1)*. Ambulatory blood pressure monitoring, the gold-standard for hypertension diagnosis, or a recently developed cuffless BP monitoring system *(42)* can be used to test the effect of circadian dosage time on efficacy. In parallel, the development of robust circadian biomarkers will help clinicians administer treatment at optimal times. In accompanying work, we report a biomarker set capable of reporting circadian phase to within 3 h from a single sample. We hope that collectively these works foster prospective model organism and clinical studies and catalyze the field of circadian medicine.

## Materials and Methods

### GTEx donor sample information

Donor inclusion criteria: 1) 21 ≤ Age (years) ≤ 70; 2) 18.5 < Body Mass Index < 35; 3) time between death and tissue collection less than 24 hours; 4) no whole blood transfusion within 48h prior to death; 5) no history of metastatic cancer; 6) no chemotherapy or radiation therapy within the 2 years prior to death; 7) unselected for presence or absence of diseases or disorders, except for potentially communicable diseases that disqualify someone to donate organs or tissues was also disqualifying for GTEx *(12)*. Additional donor sample information in Table 1.

### Preprocessing for CYCLOPS

GTEx RNA-seq used Illumina TruSeq RNA protocol, averaging ~50 million aligned reads per sample. Details at GTEx’s documentation page: https://gtexportal.org/home/documentationPage. For each tissue, gene-level TPM data was filtered to exclude any gene with a read count of zero (TPM = 0) in any sample. Following this, only the top 15,000 expressed genes by median TPM were considered for each tissue.

### CYCLOPS run parameters and analysis of transcript rhythms by cosinor regression

We used circadian seed gene inputs to sharpen CYCLOPS ordering, as reported in Anafi et. al., with the following deviations: Oscope *(43)* was used to select the eigengene clusters for ordering for each tissue (as described by Wu et. al., in the accompanying manuscript). Each eigengene cluster yielded a separate sample ordering. For the final ordering, we selected the eigengene cluster that identified rhythmic expression of clock genes and phase relationships best matching those observed in mouse. Cosinor regression was run as implemented in Anafi et. al. CYCLOPS package. Criteria for detecting rhythmic expressed genes: Entrez ID, FDR < 0.05, rAmp ≥ 0.1, RSQ ≥ 0.1.

### Statistical analyses and data visualization

CYCLOPS and cosinor regression were run in Julia version 0.3.12. All other analyses were run in R version 3.3.1 *(44)*, with packages tidyverse *(45)*, CircStats *(46)*, upsetR *(23)*, and several helper functions including drugbankR *(47)*, homologene *(48)*, ggthemes *(49)*.

### Phase set enrichment analysis (PSEA)

PSEA was run on cycling genes in human (FDR < 0.05, rAmp ≥ 0.1, RSQ ≥ 0.1) and mouse-matched (JTK Q-value ≤ 0.05) tissues to identify phase-enriched pathways from among 4653 GO:biological process gene sets from the Molecular Signatures Database.

### Drug target analyses

We used DrugBank *(25)* version 5. 0. 11, released 2017-12-20 to identify genes whose protein products are targets, transporters, carriers, or enzymes are linked to specific drug entities. DrugBank contains a total of 10,991 drug entries linked to 4,901 target protein sequences.

## Supplementary Materials

fig. S1. Pipeline for sample selection and CYCLOPS-ordering for each of 13 GTEx tissues.

fig. S2. Cycling gene counts by statistical thresholds and noncoding element type.

fig. S3. Phase set enrichment in human and mouse counterpart tissues.

fig. S4. Gene expression tracings for all cardio-cyclers that are drug targets.

## Acknowledgments

We thank Jake Hughey, Rafa Irizarry, Tiago de Andrade, and Garrett Rogers for thoughtful discussions.

## Author contributions

M.D.R., G.W., R.C.A, and J.B.H. designed research; M.D.R., G.W., and R.E.S. performed research; G.W., R.C.A., and J.B.H. contributed analytic tools; M.D.R., G.W., D.F.S., L.J.F., and R.E.S. analyzed data; M.D.R, D.F.S., L.J.F., G.W., R.E.S., and J.B.H. wrote the paper.

